# An injectable granular hydrogel stabilized by electrostatic interactions between hyaluronic acid-based microparticles and soluble gelatin exhibits poroelasticity and strain-stiffening

**DOI:** 10.1101/2025.06.09.658744

**Authors:** Julia Tumbic, Christopher B. Highley

**Affiliations:** Biomedical Engineering, University of Virginia; Chemical Engineering, University of Virginia

## Abstract

Injectable hydrogels with shear-thinning and self-healing properties are critical for biomedical applications including 3D bioprinting and regenerative medicine. While granular hydrogels inherently exhibit these properties, they often lack post-injection stability. Here, we developed an electrostatically stabilized granular hydrogel system composed of norbornene-modified hyaluronic acid (NorHA) microgels and cationic gelatin ((+) Gel). NorHA microgels (9.91 ± 4.85 μm diameter) were synthesized via batch emulsification, while (+) Gel was prepared by modifying gelatin with ethylene diamine to increase zeta potential from 2.08 ± 0.97 mV to 13.76 ± 1.11 mV. The negatively charged NorHA microgels formed stable materials when combined with (+) Gel through electrostatic interactions, confirmed by gel inversion tests and salt sensitivity studies. Rheological characterization revealed that (+) Gel addition produced poroelastic behavior and strain-stiffening properties, with storage modulus and yield onset increasing under compression. Large amplitude oscillatory shear analysis showed strain-stiffening behavior (e3 > 0) that enhanced with both (+) Gel concentration and compression. Confocal microscopy demonstrated tunable porosity through gelatin fraction control, with (+) Gel forming aggregate-like clusters. Extrusion testing showed formulations required low injection pressures (0.47-0.91 kPa) comparable to PBS and significantly lower than Pluronic, while forming robust filaments up to 23 mm in length. The materials exhibited rapid self-healing behavior and maintained structural integrity post-extrusion. This electrostatically stabilized granular hydrogel system offers a promising platform for injectable biomaterials that combine ease of delivery with post-injection stability for wound healing and 3D bioprinting applications.

## Introduction

Injectable hydrogels are important materials in a range of biomedical research applications, from 3D printing, where inks must be deposited through nozzles to create intricate structures, to regenerative medicine and therapeutic delivery for wound healing^1,2^, cancer stem cell targeting^3^, or rheumatoid arthritis treatment^4^. Two main properties of injectable hydrogels that are typically a focus of design are: 1) shear-thinning behavior, and 2) self-healing behavior. To extrude through a needle, hydrogel structure must be disrupted to allow material flow. In self-healing, where the structure of a hydrogel material must be reestablished, with the material designed to recover from the disruption and reform a gel mechanical properties similar to the original. To achieve these properties within continuous hydrogels, dynamic bonds within the polymeric network, including hydrazone bonds^5,6^, guest-host bonds^7^, and electrostatic interactions^6,8,9^, have engineered since these bonds easily brake and reform upon application and removal of a force. Shear-thinning behavior is also inherent in some materials, such as hyaluronic acid^10^. Self-healing has also been engineered or supported by designing interpenetrating networks (IPNs)^11,12^, semi-IPNs^13^, and embedding nanoparticles into hydrogels, where one of the components in these multicomponent systems can aid in restoring mechanics post-damage^14,15^.

Granular materials, present inherent shear-thinning and self-healing behaviors, and thus are attractive as injectable and extrudable materials^16^ in applications such as minimally-invasive deliver of regenerative biomaterials^1^ or 3D bioprinting^17^. Granular systems are comprised of particles rather than a bulk gel, and when particle rearrangements occur at sufficiently high forces^18^ they can be easily be extruded through small nozzles. In comparison to continuous hydrogel systems, where high forces permanent disrupt the original structure, granular hydrogels flow upon application of force and revert to a stable solid-like state when force is removed, recovering mechanical properties from the pre-extruded state. However, absent any designed interparticle interactions, granular hydrogels may be weak as bulk materials after deposition: they may flow again if sufficient force is applied and are subject to surface erosion of constituent particles. Covalent crosslinking^1^ (or annealing) between particle surfaces or non-covalent interactions, including guest-host bonds^19^, hydrazone bonds^20^, and electrostatic interactions^9,21–23^ can be used to stabilize granular materials.

Here, we describe the design of an injectable, electrostatically stabilized granular hydrogel formed from biomaterials commonly used in regenerative medicine and tissue engineering applications, hyaluronic acid (HA) and gelatin. Because HA is an anionic polymer polymer, positively charged materials will associate with it via electrostatic interactions. Here, a norbornene modification was introduced on the backbone of the HA (NorHA) to allow microgels to be stabilized from batch emulsions through full consumption of the norbornene functionalities. Gelatin was used to generate a positively charged material that could electrostatically bind particles to one another. To create a cationic polymer, (+) Gel, that could adsorb onto the NorHA microgels, we modified gelatin type A with additional amine groups. In characterizing this new material system, we first studied the interactions between the NorHA particles with (+) Gel, and the effects of packing fraction and (+) Gel concentration on these interactions. Our characterization showed that the addition of (+) Gel produced a granular material with poroelastic-like effects as well as strain-stiffening behavior. We then examined a formulation with varying (+) Gel concentrations on the potential use of this material as an injectable system. We saw that (+) Gel adhered particles together to form filaments even at low concentrations but were still easily extrudable. This material offers an approach to designing injectable biomaterials that inherently stabilize upon injection. It also presents a platform with poroelastic and strain-stiffening behaviors. The former quality is observed in tissues like cartilage^24,25^ and the dermis^26^ and is important in pathologies like as fibrosis^27,28^, and the latter is characteristic of several native biopolymers^29^ but is must be engineered into synthetic systems where controlled mechanical properties are desired^30–34^.

## Methods

### Materials synthesis

Norbornene-modified hyaluronic acid (NorHA) was prepared as described in chapter 1, using a previously established method^35^. Briefly, HA was dissolved in DI water to achieve a 2wt% solution, followed by addition of an ion exchange resin (Dowex). This reaction was then filtered over a Buchner funnel and the pH adjusted to roughly 7 prior to lyophilization. The synthesized HA-TBA was dissolved in anhydrous dimethylsulfoxide (DMSO) and reacted with 5-norbornene-2-carboxylic acid and 4-(dimethylamino)pyridine (DMAP), catalyzed with di-tert-butyl decarbonate (BOC_2_O). The reaction was run overnight. The resulting material was dialyzed against DI water for three days, followed by purification via ethanol precipitation, resuspension in DI water, and further dialysis against DI water for an additional three days. This material was lyophilized and stored at -20°C until needed. The degree of modification was characterized via H^1^ NMR in D_2_O.

Cationic gelatin ((+) Gel) was prepared using a modified published procedure^36^ by reacting gelatin Type A (300 bloom) at 2% in PBS with 1-ethyl-3-(3-dimethylaminopropyl)carbodiimide (EDC HCl, Sigma), N-hydroxyl succinimide (NHS, Sigma), and ethylene diamine (EDA, Sigma) (Fig. 1B). Per gram of gelatin, the weight ratios of reactants were 1:0.6:3.6 of EDC:NHS:EDA. The gelatin was first dissolved in PBS at 37°C, then NHS and EDA were added to the solution at room temperature. The pH was adjusted to 5.5 to 6, after which the EDC was added. The pH was then further adjusted to 4.5 to 5 and allowed to react for approximately 24h at room temperature. The (+) Gel was then dialyzed against DI water for 2-3 days at room temperature. A fluorescamine assay (Sigma) was used to determine the change in amine groups using a calibration curve with glycine and compared with unmodified gelatin. The assay protocol was modified from a previously published study^37^, where the fluorescamine powder was first dissolved in DMSO at a concentration of 3mg/mL. Gelatin solutions were diluted to roughly 1.25mg/mLin PBS but adjusted depending on fluorescence values. Then, 30µL of assay was added and mixed with 90µL of gelatin solution. The assay and gelatin were allowed to react for 15 min prior to running fluorescence testing.

**Fig. 1:**
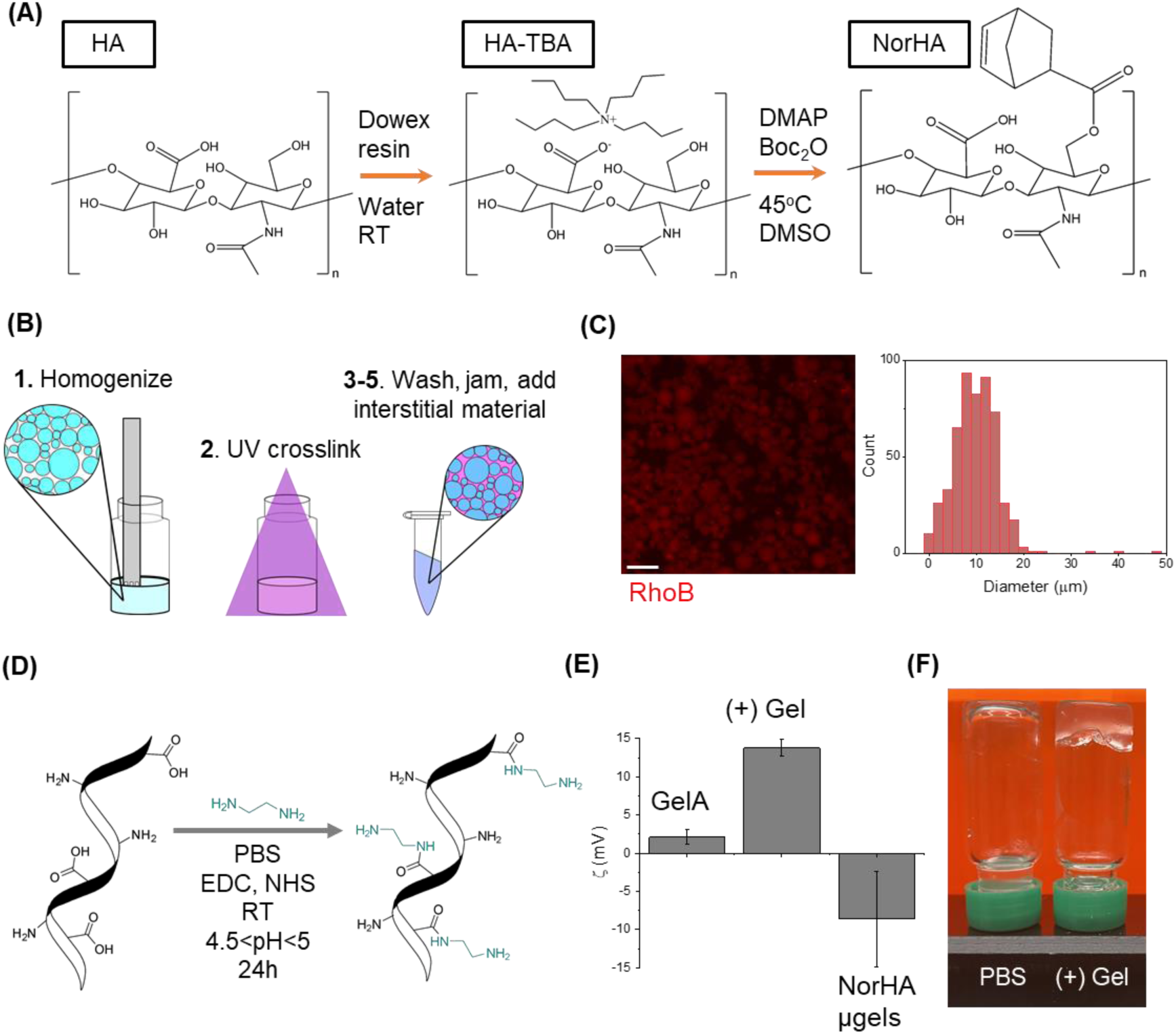
(A) NorHA synthesis route, (B) NorHA microgel fabrication, (C) widefield image of NorHA microgels (scalebar=40um) with particle size distribution (n=524), (D) (+) Gel synthesis route, (E) zeta potential of gelatin type A, (+) Gel, and NorHA microgels (n=3, * represents p<0.05, and error bars denote standard deviation) and (F) photo of particles at dilute packing densities with PBS or (+) Gel, illustrating how the (+) Gel can tether particles together to form a gel-like material.

**Fig. 2:**
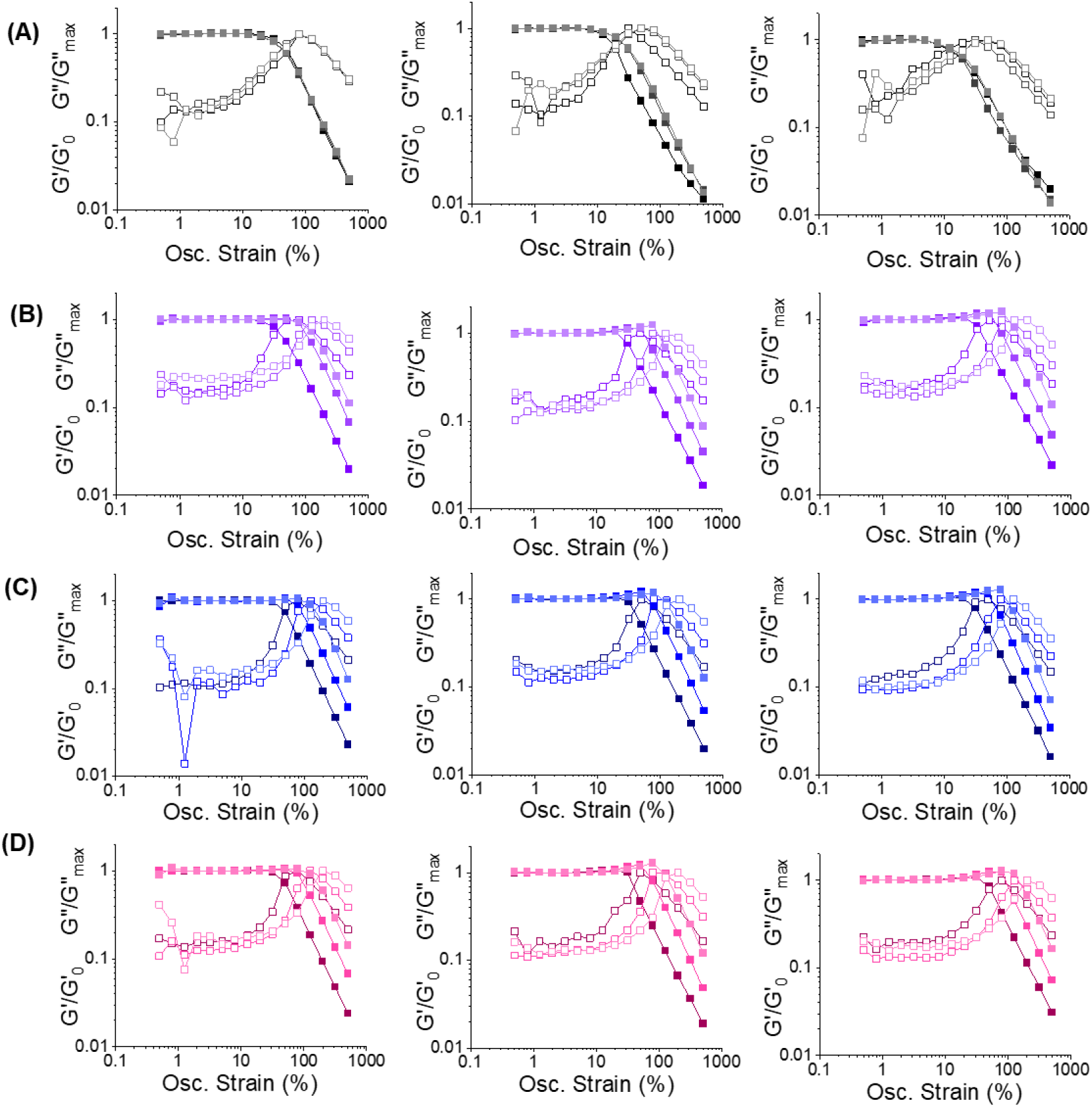
Normalized strain sweeps at different compressive strains (shown by color gradients where lighter denotes increased compressive strain), jamming fraction (going across row) and (+) Gel concentration **A)** black=0 µg/mg, **B)** purple=3 µg/mg, **C)** blue=6 µg/mg (5 µg/mg for 0.5 jamming fraction), and **D)** pink=9 µg/mg (10 µg/mg for 0.5 jamming fraction)). Storage moduli were normalized to G0 from the linear viscoelastic regime, and loss moduli are normalized against the maximum value found in the original loss moduli data.

### Granular system preparation

#### NorHA microgel fabrication

A precursor 3 wt% NorHA solution was prepared in PBS containing 0.313 mg/mL dithiothreitol (DTT, Sigma) and 6.6mM lithium phenyl-2,4,6-trimethylbenzoylphosphinate (LAP, Sigma). This NorHA solution was homogenized in light mineral oil with 2% Span80 for 2 minutes at 3000rpm, followed by UV curing for 5min at approximately 10mW/cm^2^. De-emulsification was started by adding isopropanol to the emulsion post-crosslinking. After stirring, the mixture was filtered over a 0.22um hydrophilic PVDF membrane. Microgels were washed an additional two times over the filter with isopropanol. After filtering, the microgels were left to dry then added into a 50mL conical and vortexed with 70% ethanol and placed on a rocker for two days. To finish processing, microgels were washed with PBS four times using centrifugation to ensure ethanol removal and equilibration in PBS. Microgels were then filtered over a 40um cell strainer and stored at 4°C until needed.

#### Jamming and interstitial material addition

First, particles were jammed according to particle weight, and the resulting volume of gelatin that was added relative to the particle weight is defined as gelatin fraction in this study. Particles were jammed in pre-weighed eppendorf tubes at 21000xg for 5 minutes. Excess PBS or media was then removed. Particles were centrifuged again if needed to fully remove excess PBS or media. Once particles were packed, the particles were weighed. Based on this weight, gelatin solutions were added to achieve a 0.3 gelatin fraction. To achieve a 0.0 gelatin fraction, the particles were centrifuged again at 21000xg and excess solution was removed. To achieve a 0.5 gelatin fraction, additional PBS was added to the particles corresponding to the original weight of the particles.

### Zeta potential measurements

All materials were prepared in filtered 0.1X PBS. For the gelatin samples, all samples were prepared at 5mg/mL. NorHA microgels were tested at 3mg/mL due to their size^38^. Zeta potentials were measured using a zetasizer (Zetasizer Ultra, Malvern) and calculated using the Smoluchowski equation. Each sample was run three times with independently-prepared samples.

### Confocal imaging of interstitial space, and porosity and pore size measurements

NorHA microgel scaffolds were prepared at different gelatin fractions and (+) Gel concentrations. The interstitial gelatin contained high molecular weight FITC-dextran (2MDa) to aid in visualization during image acquisition. Images were acquired using a confocal microscope (Stellaris 5, Leica). Z-stacks were taken with slices set to 2µm for a total depth of 25-40µm. Area fraction was calculated from three independently-prepared samples for each gelatin fraction and (+) Gel concentration using FIJI. A 2D slice was thresholded then the measure function was used to obtain area fraction. For each group, three independently-prepared samples were used.

Average pore size and pore size distributions were estimated using a FIJI plug-in, BoneJ. Z-stacks obtained from confocal microscopy were first resliced to obtain isotropic pixels. The resliced stack was then thresholded using Otsu’s method^39,40^. The thickness function in BoneJ^41^ was used to calculate trabecular thickness, which returns the average pore size within the scaffold. The resulting thickness map that is also calculated from this function was used to obtain pore size distributions (SI Fig. 7). An n of 2-3 independently-prepared samples was used for each group.

Porosity and pore size were measured for samples compressed to 30% and 60%. To achieve these compressions, custom-made plugs with thicknesses of 1mm or 2mm were printed using a PLA printer (Prusa) that would achieve 33% and 67% compressions if the material were placed in a PDMS holder of 3mm (SI Fig. 8). The materials were set up as described above and compressed right before imaging on a confocal microscope. Z-stacks of similar dimensions were obtained and used to quantify area fraction and pore size. Measurements were done in duplicate in independently prepared samples.

As described above, inks containing 0-3% (+) Gel with high molecular weight FITC-dextran were extruded at flow rates of 11.68mL/h and 1.17mL/h into PDMS holders plasma-bonded to glass-bottom 6-well plates. Light mineral oil was placed on top to prevent dehydration during imaging. Imaging was done to obtain z-stacks that were then used to calculate porosity and pore size using FIJI as described above. Measurements were done in triplicate with three independently-extruded samples.

### Rheology

NorHA particles were jammed with 0-3% (+) Gel at different gelatin fractions. Samples were prepared right before testing. An 8mm sand-blasted parallel plate was used in combination with a Peltier plate with sandpaper placed onto it to prevent wall slip. Testing was done at 20°C. For each sample, the top geometry was placed carefully onto the material to not compress the material, then the test started. Strain sweeps were conducted in shear from 0.5% to 500% with a frequency of 0.1Hz. After a strain sweep, the material was compressed to roughly 30%. This was followed by another strain sweep. The material was then further compressed to roughly 60% of the original sample height, and a final strain sweep was conducted. After each compression, the material was carefully trimmed prior to the start of the next strain sweep. Analysis of the change in storage modulus, critical strain (defined as departure linearity in the strain sweep), yield strain, and modulus crossover changes was carried out.

### Strain-stiffening analysis

Strain-stiffening behavior was quantified using MITLAOS software (LAOS = large amplitude oscillatory shear)^42^. Stress and strain data were taken from the original rheology files at the highest point in G’ in the strain sweep prior to yielding. To evaluate strain-stiffening behavior, the Chebyshev coefficient, e3, calculated from MITLAOS, was plotted against (+) Gel concentration without compression and as a function of compressed strain for each sample.

### Extrusion testing

A syringe holder designed to hold a 1mL syringe was 3D printed using an STL printer (Formlabs) (SI Fig. 11). Microgel formulations were prepared and added into 1mL syringes with air bubbles removed via needle, stored at 4°C and tested the next day. A 100N load cell was used on an Instron machine set up for compression testing. 25G blunt needles were added to the syringes prior to testing. The syringes were compressed at a rate of 0.5mm/s, corresponding to a volumetric flow rate of approximately 27.54 mL/h. Materials were compared to extruded PBS and 30% Pluronic. The change in pressure over volume was then calculated and graphed.

Extruded filaments for each ink were examined visually. For each ink, 50µL was extruded at either a high flow rate of 11.68mL/h or a low flow rate of 1.17mL/h. Each extrusion was repeated five times. There was a pause of at least two minutes between extrusions. Videos were acquired using an iPhone 11, and filament lengths were measured using FIJI.

### Statistical analysis

Statistical analyses were carried out using Origin. One-way ANOVAs were used with a post-hoc Tukey test to determine statistical significance and specify differences between groups. A p-value of less than 0.05 was considered statistically significant.

## Results and Discussion

### Materials characterization

Norbornene-modified hyaluronic acid (NorHA) (Fig. 1A) was prepared with between 16% and 21% modification of the HA disaccharide unit (SI Fig. 1). NorHA microgels formed via batch emulsification (Fig. 1B) had average particle diameters of 9.91 +/- 4.85 µm (Fig. 1C and 1D). The particle size distribution was broad, which was expected due to the method used to form the particles. This was confirmed by calculating the coefficient of variation, which was 48.99%.

Cationic gelatin ((+) Gel) introduced amine groups at carboxylic acids already present on the gelatin backbone (Fig. 1D). By doing so, negatively charged groups were eliminated and replaced with positively charged groups, increasing the zeta potential of the gelatin markedly from 2.08 +/- 0.97 mV to 13.76 +/- 1.11 mV (Fig. 1E). This modification was confirmed further with a fluorescamine assay, which showed an increase from 224.65 +/- 21.54 µmol NH_2_ per g material to 500.08 ± 35.76 µmol NH_2_ per g material (SI Fig. 3).

**Fig. 3:**
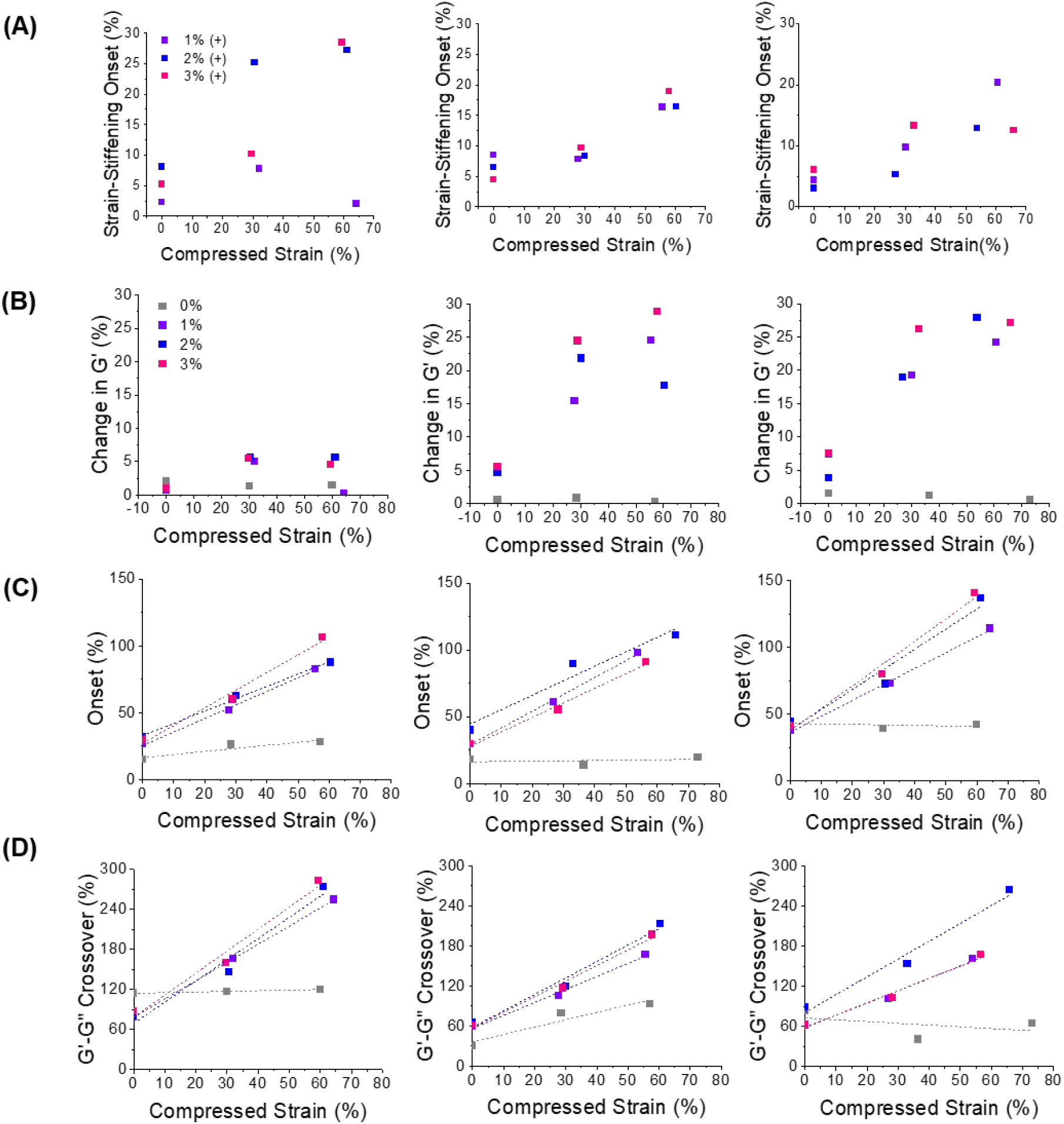
Change in (A) critical strain, (B) storage modulus, (C) yielding onset, and (D) moduli crossover as a function of (+) Gel concentration and compressive strain, at (i) 0, (ii) 0.3, and (iii) 0.5 gelatin fractions. For granular materials tested at a gelatin fraction of 0.5, (+) Gel concentrations of 0% (gray), 0.6% (purple), 1% (blue), and 2% (pink) were used.

**Fig. 4:**
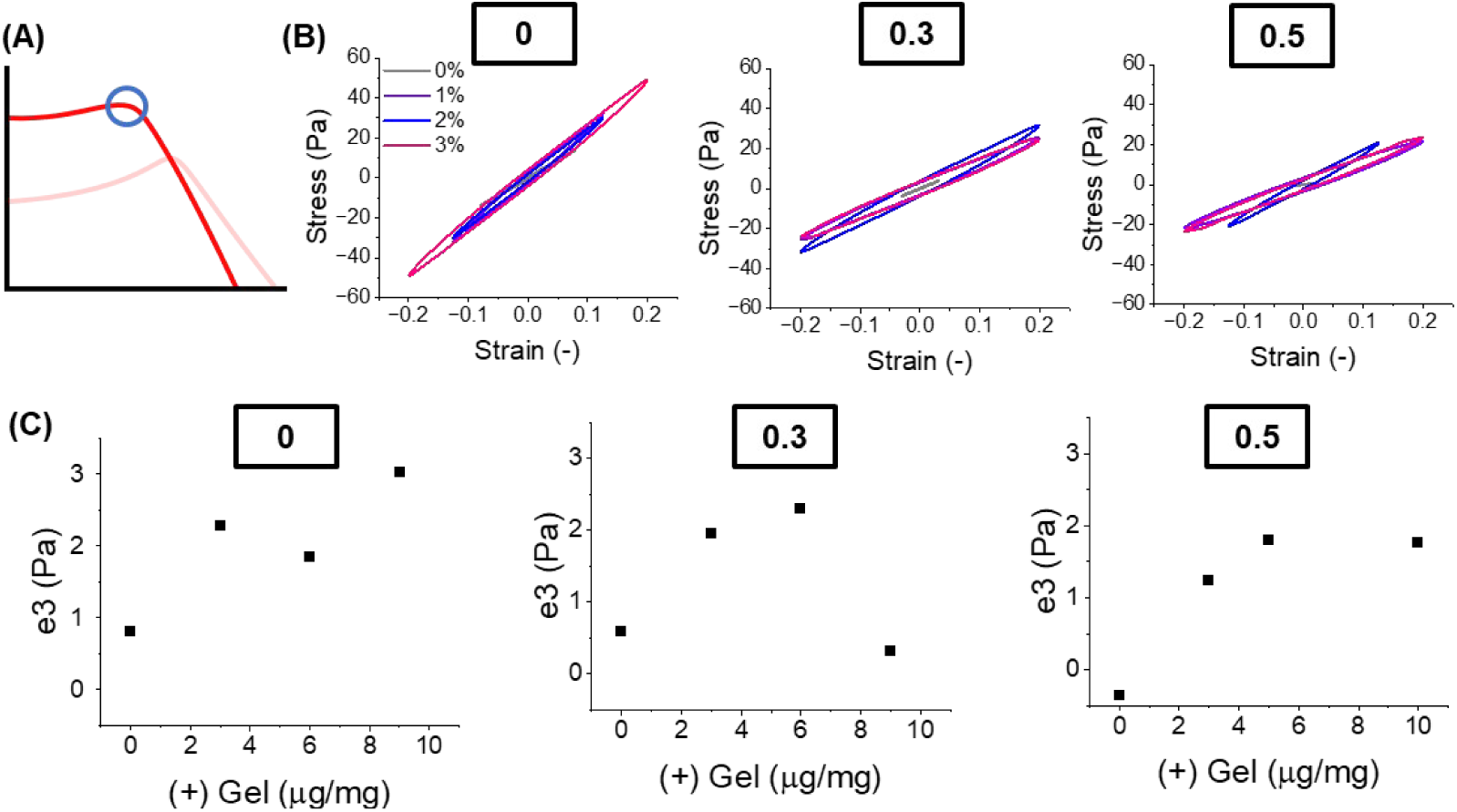
Large amplitude oscillatory strain analysis (A) diagram of where point in strain sweep is being used in LAOS analysis, (B) LAOS at different (+) Gel concentrations and gelatin fractions under no compression, and (C) change in the Chebyshev coefficient, e3, as a function of (+) Gel concentration and gelatin fraction.

The zeta potential of the NorHA microgels was -8.64 ± 6.23 mV. When added to the NorHA particles, the (+) Gel appeared to tether the particles together, keeping it in a gel-like state even at dilute packing densities compared to PBS alone. This could readily be seen in gel inversion tests (Fig. 1F), where the NorHA particles surrounded by PBS flowed from the base of a vial to the top when the vial is inverted, while particles surrounded by (+) Gel did not flow and remained in the base of the vial, a behavior that is characteristic of a solid hydrogel.

### Rheology

During rheological testing, we observed in preliminary tests that formulations of NorHA with (+) Gel ejected PBS as the top geometry compressed the sample. This is indicative of poroelastic behavior^43^. To test this further, oscillatory strain sweeps were conducted at compressive strains of 0%, 30% and 60%. A similar test was used previously for studying poroelastic polymer networks and was adapted for this study^43,44^, with the difference that the granular system here derives bulk mechanics from particle interactions. It was expected that as (+) Gel-containing granular hydrogels were compressed, interactions between the NorHA microgels and the (+) Gel that adsorbed onto the particles would increase, leading to changes in viscoelastic properties, such as yielding behavior. Nonetheless, despite increased interactions, the particles were still expected to be able to rearrange since the interactions are non-covalent. In contast, in a bare particle system that did not contain (+) Gel, we expected that the particles would freely rearrange during compression and exhibit no changes in the rheological results with compressive strain.

Gelatin fraction and (+) Gel concentration were varied to assess the influence of (+) Gel and porosity on yielding behavior of these formulations as a function of compressive strain (Fig. 2 and SI Fig. 3). For formulations containing no (+) Gel (Fig. 2A), the strain sweeps at each compressive strain were nearly identical at all tested gelatin fractions. This indicates that particle rearrangements freely occur as the material is compressed, as expected, and therefore poroelastic behavior is not a feature of PBS-based materials. Once (+) Gel is introduced (3 µg/mg in Fig. 2B, 6 µg/mg in Fig. 2C, and 9 µg/mL in Fig. 2D, except as noted), critical strain, storage modulus, yielding onset, and modulus crossover values generally increase as the material is compressed at any (+) Gel concentration. These values are extracted from the rheological measurements and presented in Fig. 3. These data indicate that when (+) Gel was introduced, particle rearrangements became restricted by electrostatic binding in these formulations and, therefore, that the (+) Gel acted enhance interparticle interactions through increasing particle density as water was removed from the system. However, because electrostatic interactions are dynamic in nature, particle rearrangements still occurred. Interestingly, (+) Gel concentration appeared to not influence bulk storage modulus when no compressive strain is present. At all gelatin fractions, bulk storage moduli were the same (SI Fig. 5), with comparison to PBS-based formulations having moduli values that were generally smaller. It is possible that this could be a function of bulk rheology testing specifically, where at small strains and frequencies, the differences in (+) Gel concentration may not be apparent.

**Fig. 5:**
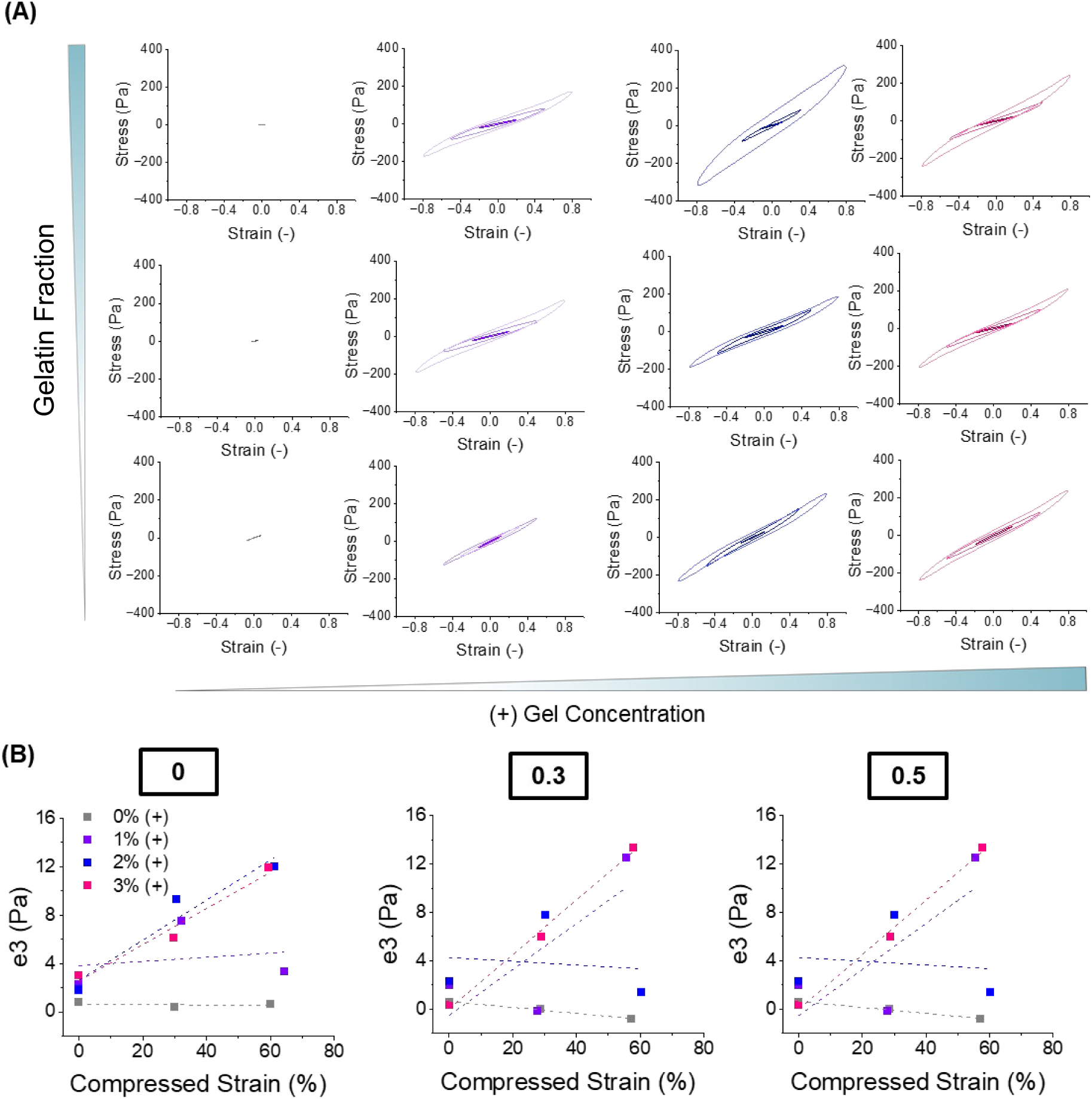
LAOS sweeps performed under compression show (A) Lissajous curves assuming sigmoidal shapes indicative of strain stiffening as functions of gelatin fraction and (+) Gel concentration, and (B) there is a corresponding change in the Chebyshev coefficient, e3, indicating strain stiffening as a function of compressive strain, (+) Gel concentration, and gelatin fraction.

To confirm that rheological measurements were a function of the (+) Gel chains and the NorHA particles interacting electrostatically, rheology of these systems with varying salt concentrations and 1% and 2% Tween20 were analyzed (SI Fig. 6). An increase in salt concentration leads to shielding of charges and allows for testing for effects of electrostatic interactions, whereas the addition of Tween20 interferes with hydrophobic interactions that might be introduced with (+) Gel^45^. Strain sweeps showed little to no change in the systems containing either 1% or 2% Tween20, indicating that there are little to no hydrophobic interactions contributing to the bulk rheological properties (SI Fig. 6B). The addition of salt to the system, however, caused a decrease in the storage modulus, yield onset, and modulus crossover. This confirms that electrostatic interactions play a role in the bulk rheological properties (SI Fig. 6A). This can also be seen visually, where a sample containing 1% Tween20 appeared to retain the gel-like properties seen in the non-treated particulate system. This contrasts with the appearance of the gel when salt is added. In the latter case, there is a clear difference where the jammed system becomes increasingly liquid-like (SI Fig. 6C).

**Fig. 6:**
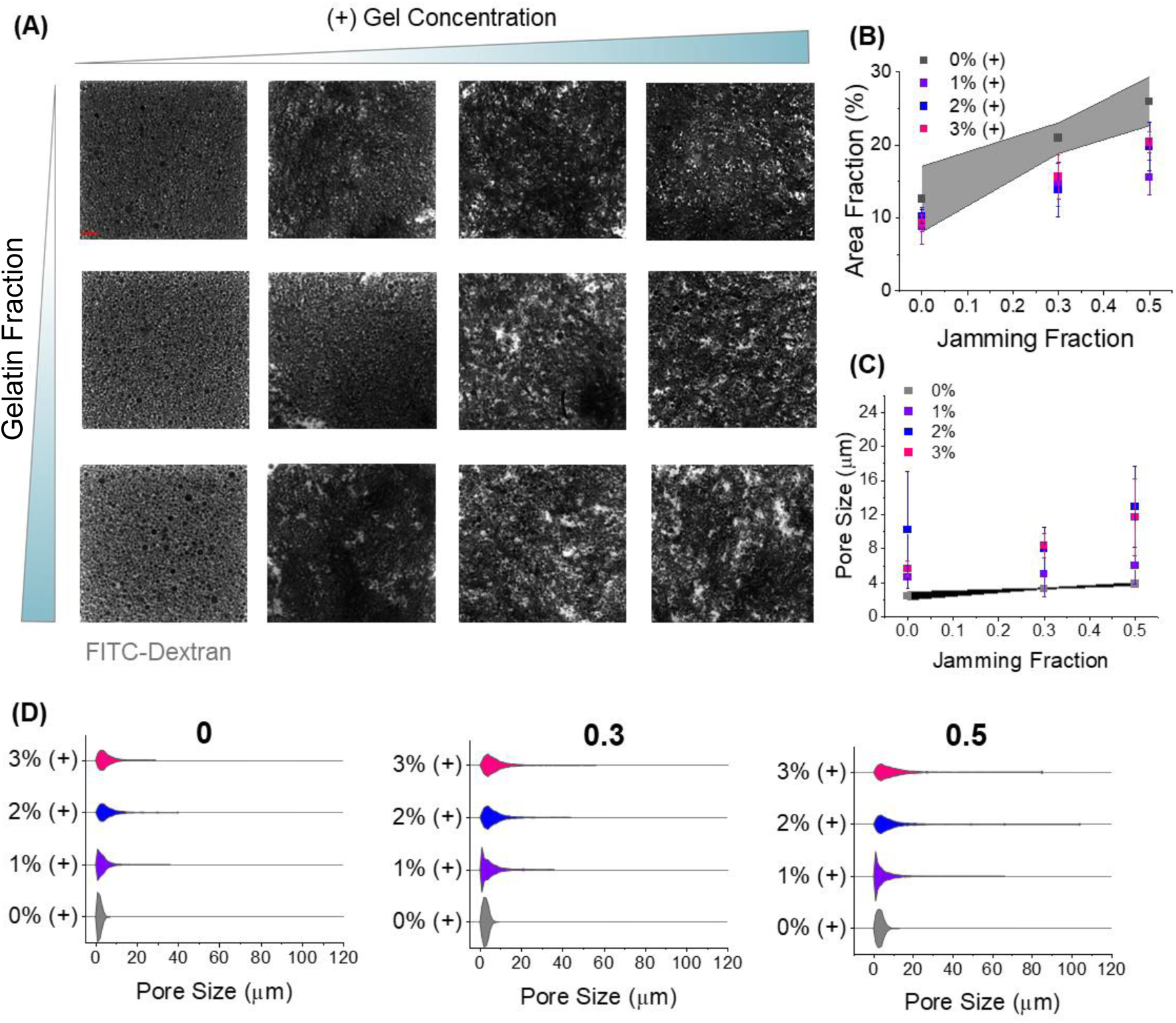
Microporosity within the granular system (A) increases with gelatin concentration and gelatin fraction as seen in confocal images, and quantified in (B) change in area fraction as a result of gelatin fraction and (+) Gel concentration, (C) change in average pore size as a result of gelatin fraction, and (D) pore size distributions due to changes in (+) Gel concentrations and gelatin fractions.

**Fig. 7:**
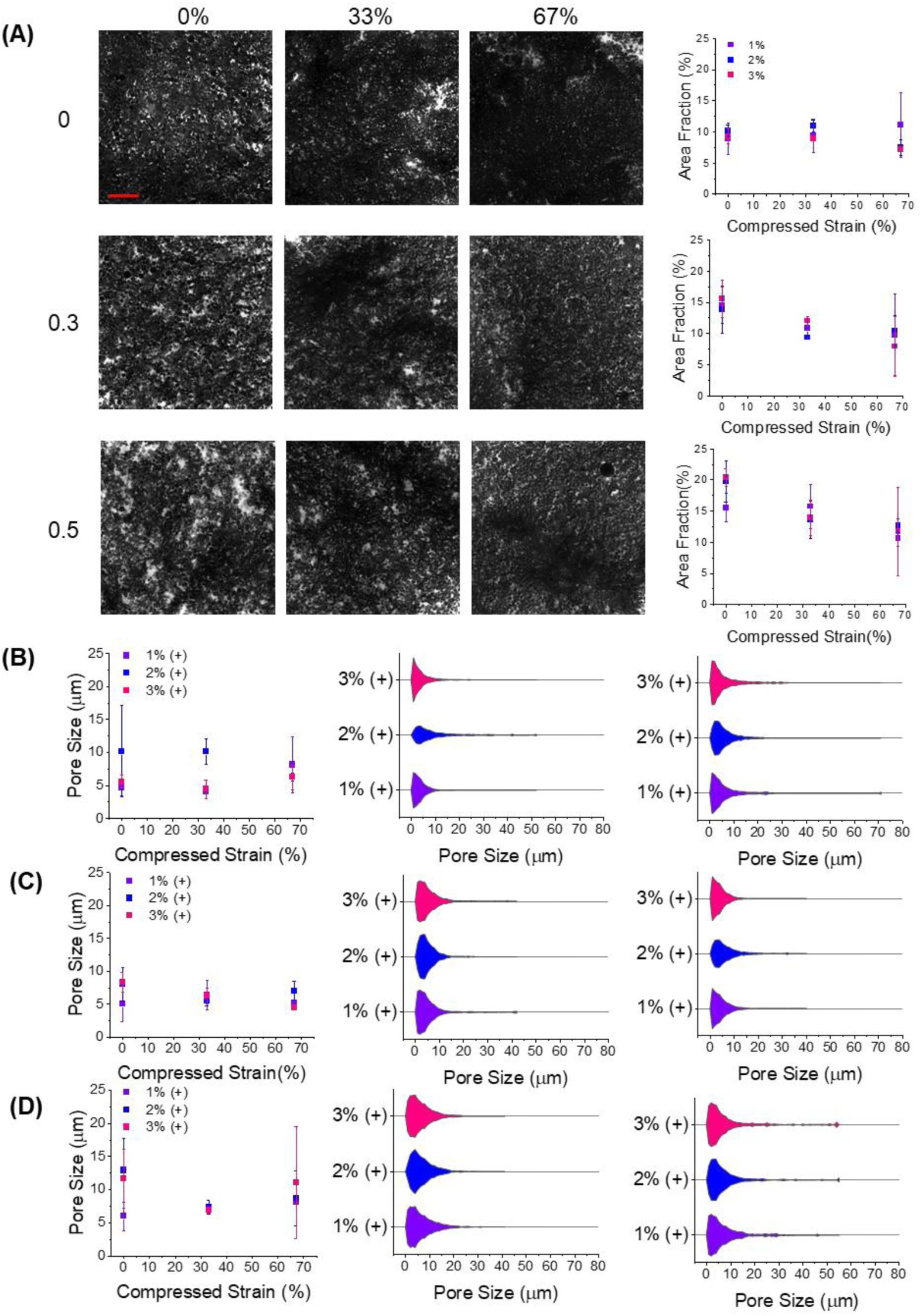
Porosity decreases with compression and increasing gelatin fraction. (A) Confocal images of 3% (+) Gel formulations compressed at 0%, 33%, and 67% strains at 0, 0.3, and 0.5 gelatin fractions and with the corresponding graphs showing change in area fractions, and – for gelatin fractions of (B) 0, (C) 0.3, and (D) 0.5 – changes in (i) average pore sizes and (ii) pore size distributions (2 samples shown).

**Fig. 8:**
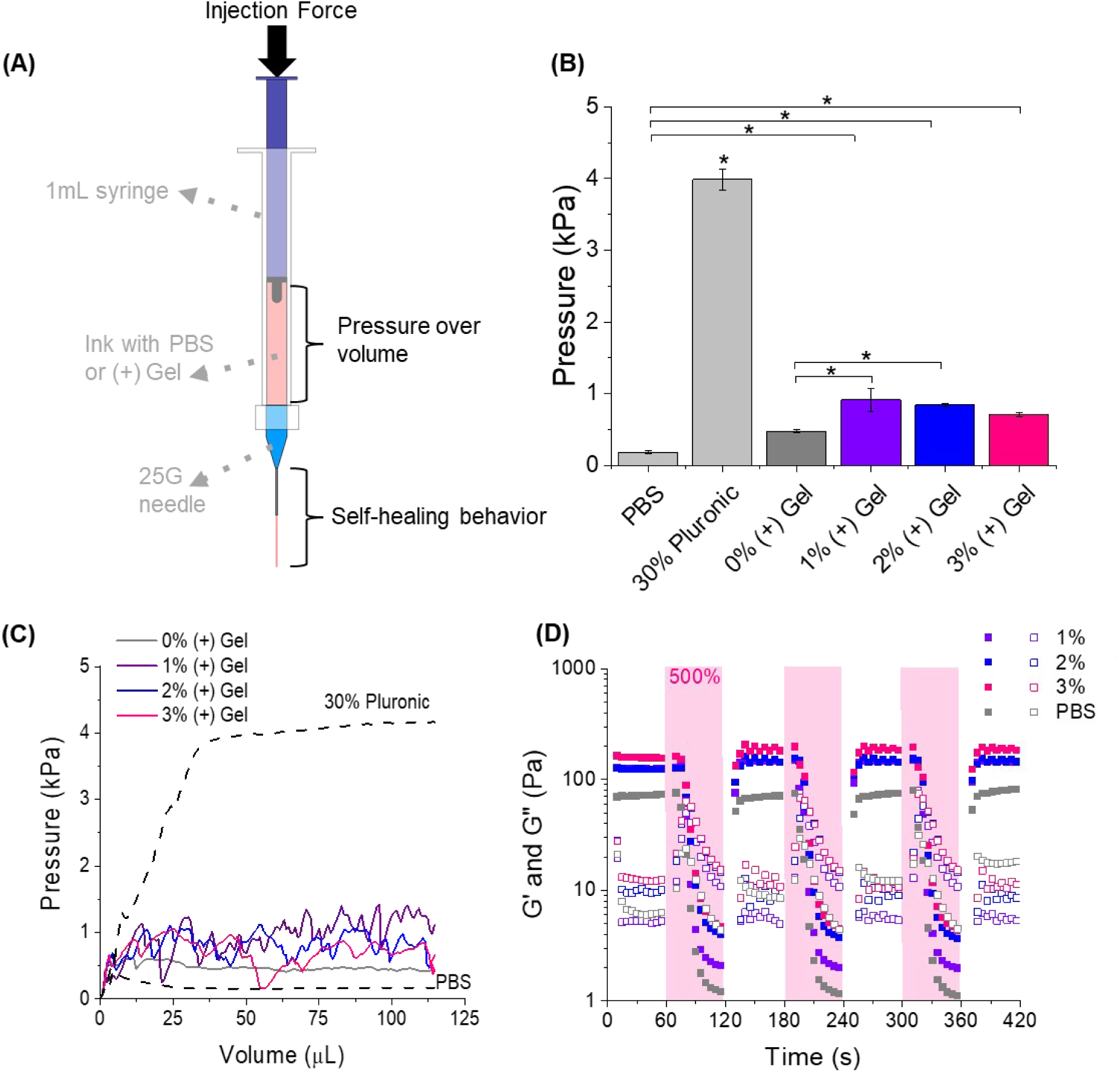
(A) Schematic of variables tested for assessing injectability of formulations with different (+) Gel concentrations, (B) average pressure during extrusion with Pluronic F-127 and PBS used for comparison (n=2, * denotes p<0.05, and error bars represent standard deviation), (C) representative data of pressure during extrusion (x-axis is total volume extruded, a proxy for time) of formulations as a function of (+) Gel concentrations,with Pluronic and PBS for comparison, and (D) repeated high and low strain rheology showed self-healing behavior of the materials over several ‘injection’ cycles as a function of (+) Gel concentration.

In addition to linear increases in the onset and moduli crossover points, there was a noticeable strain-stiffening behavior in the formulations containing (+) Gel. To analyze this behavior, Lissajous curves were generated at the points of strain-stiffening in large amplitude oscillatory strain (LAOS) sweeps. These curves capture the changes in behavior with (+) Gel concentration and applied compressive strains. For comparison, the points prior to yielding in the PBS-based formulations were used (Fig. 4A). Under no compression, all materials showed primarily viscoelastic behavior, as expected, where the Lissajous curves are linear but open (Fig. 4B). As compression is introduced, a noticeable change in the shape of the Lissajous curves for (+) Gel-containing formulations is observed. With compression, the Lissajous curves take on an inverted sigmoidal shape, indicative of strain-stiffening behavior (Fig. 5A). This was not observed in the PBS controls, as expected.

To assess nonlinear elastic behaviors in the material indicative of poroelasticity and strain stiffening, further analysis was done examining the Chebyshev coefficient, e3, for each material as a function of (+) Gel concentration. This analysis was performed both under no compression and under compressive strains. This Chebyshev coefficient is calculated via decomposition of the Lissajous curve^46^: when e3 is greater than 0, strain-stiffening behavior is present; when e3 is negative the material is strain-softening^46^. We observed that as (+) Gel concentration increased, the e3 value also increased across all gelatin fractions, suggesting that (+) Gel contributed to strain-stiffening behavior (Fig. 4C). When compression was introduced, a similar trend was seen. As formulations containing (+) Gel were compressed further, the strain-stiffening behavior was enhanced, as seen in the increase in e3 (Fig. 5B). This indicated the presence of poroelastic-like behaviors as (+) Gel-particle interactions were enhanced under compression.

### Porosity and effect of (+) Gel concentration and compression

Because (+) Gel was expected to adsorb onto the NorHA microgels, the change in particle-particle interactions was expected to cause microstructural changes of the packed microgel materials. Thus, area fractions, average pore sizes, and pore size distributions of the packed materials were assessed using confocal microscopy. For all materials, FITC-dextran, a neutrally-charged polymer, was added to the (+) Gel or PBS solutions to visualize the interstitial space (Fig. 6A). Computational tools (FIJI) were then used to quantify area fraction from the acquired images. As expected, area fraction increased with gelatin fraction, showing that the porosity within the jammed systems was easily tunable through this design parameter (Fig. 6B). Average pore size and pore size distributions were measured and assessed using an ImageJ plugin, BoneJ (SI Fig. 7). Average pore sizes in the (+) Gel-containing formulations were generally greater across all gelatin fractions compared to PBS-based formulations (Fig. 6C). This can be attributed to the difference in pore size distributions, where higher distributions were seen in the (+) Gel-containing formulations (Fig. 6D). This is due to the presence of aggregate-like clusters that seem to form when adding (+) Gel to the microgels. This type of structure has been seen previously in particle-based systems where attractive interparticle interactions were dominant^47,48^.

From the rheology data, there was a clear trend in formulations containing (+) Gel of compressive strain with respect to rheological measurements. Since this material has rheological behavior pointing to poroelastic characteristics, area fractions and pore sizes of compressed formulations containing (+) Gel were assessed. The formulations were again imaged with HMW FITC-dextran in the interstitial space (Fig. 7A and SI Fig. 9-10). Formulations were compressed to 33% or 67% using a custom-made device described in the Methods section (SI Fig. 8). Overall, a decrease in area fraction was observed as compression was increased, and this decrease seemed dependent on gelatin fraction, as higher packed materials were less affected by compression (Fig. 7A). When examining pore size distributions, there was a general shift towards smaller pore sizes as compressive strain increased (Fig. 7B-D). These results combined with rheological data indicate that these materials had poroelastic characteristics and became denser under a compressive force, that drove enhanced electrostatic interactions and increased yielding onset and moduli crossover points.

**Fig. 9:**
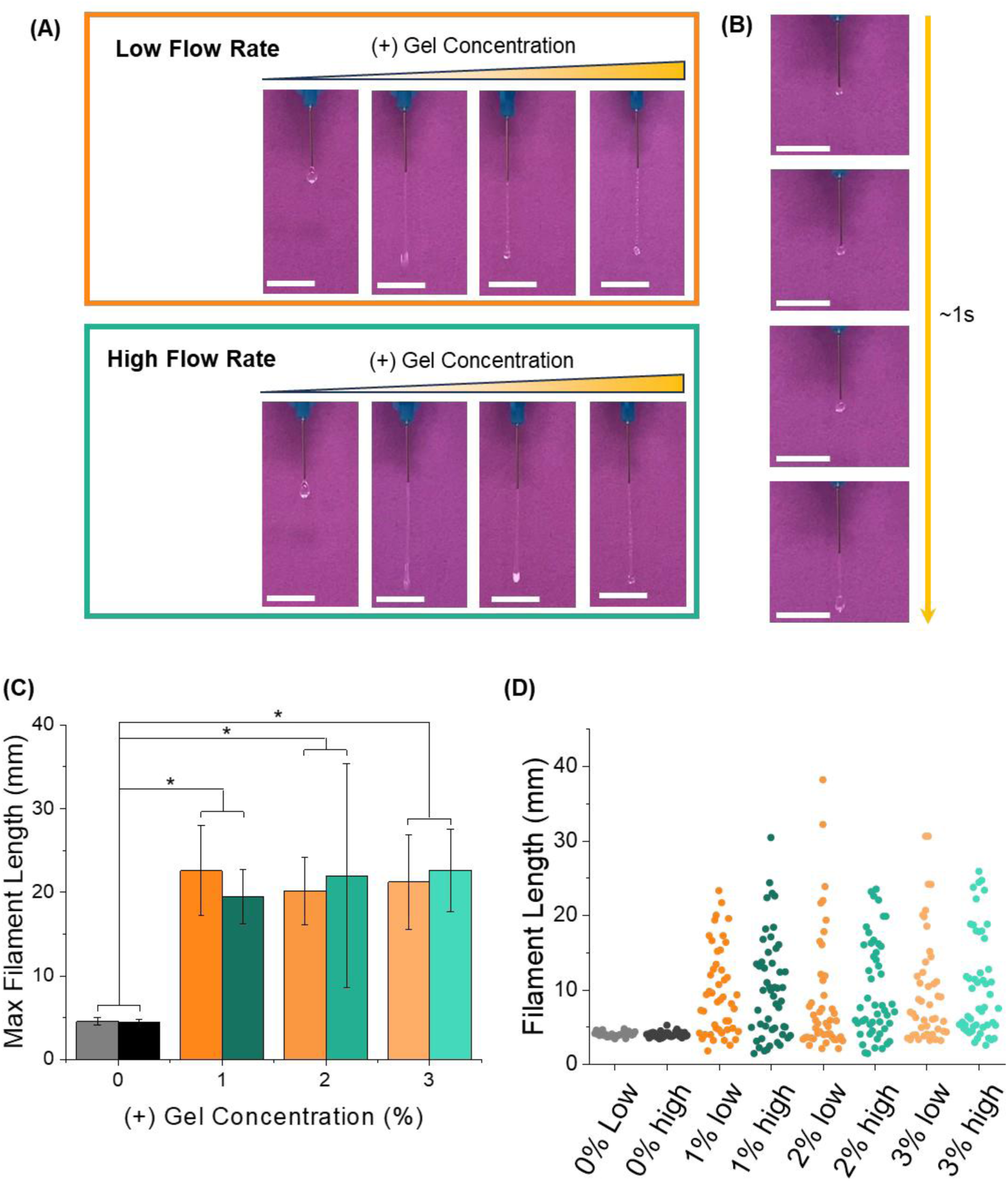
Qualities of filaments of materials extruded from a syringe needle. (A) Images of formulations containing varying concentrations of (+) Gel at low (1.2 mL/h) and high (11.7 mL/h) flow rates; (B) An example of a (+) Gel containing formulation being extruded and the material balling up during the first 1s of extrusion forming the characteristic bead of material at the end of the (+) Gel containing formulations; (C) Maximum filament lengths achieved for each extruded formulation at high (green) and low (orange) flow rates, n=5, error bars indicate standard deviation and * represents a p-value less than 0.05; (D) Distribution of filament lengths for each formulation at high (green) and low (orange) flow rates.

**Fig. 10:**
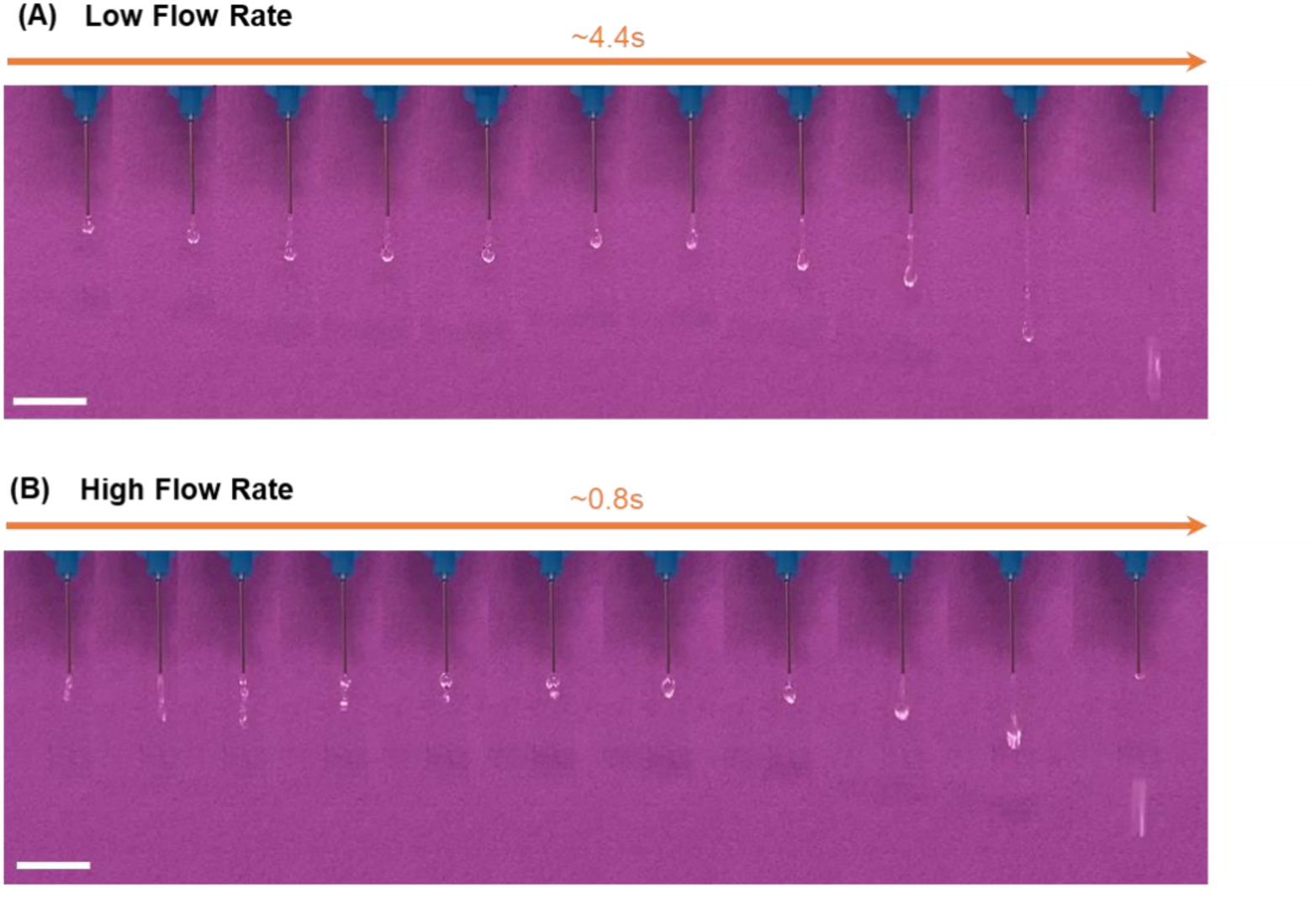
Time-lapse images of 1% (+) Gel formulations extruded at (A) low and (B) high flow rates (scalebar =1 cm).

**Fig. 11:**
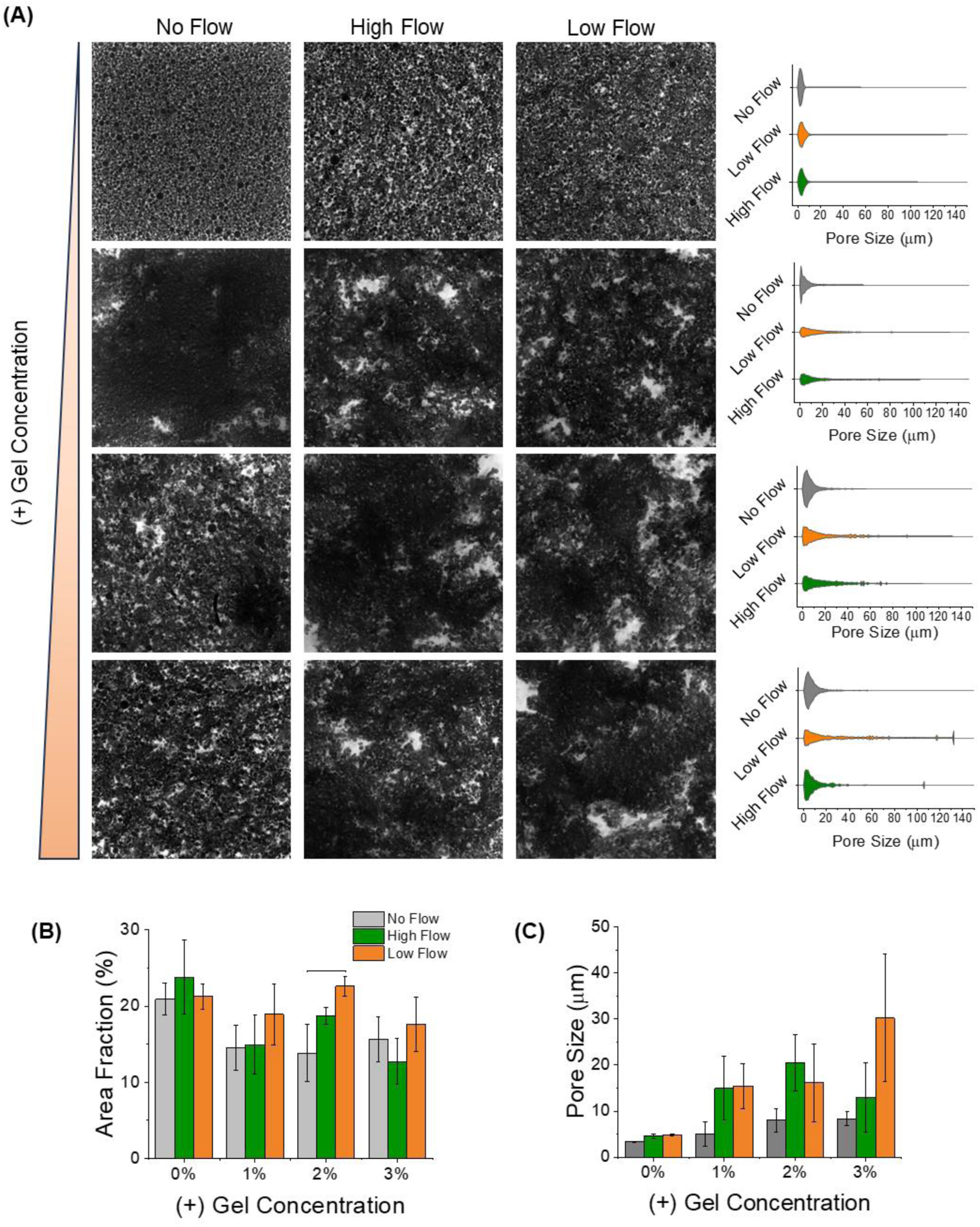
Comparing granular material microstructure before and after extrusion. (A) Confocal images of formulations with different (+) Gel concentrations under no flow, high flow, and low flow conditions with corresponding pore size distributions, (B) change in area fraction as a function of flow rate and (+) Gel concentrations (n=3, * represents p<0.05, and error bars denote standard deviation), and (C) change in average pore size as a result of flow for each (+) Gel concentration (n=3, * represents p<0.05, and error bars denote standard deviation).

### Extrusion testing and filament measurements, and porosity calculations

Three primary measurements were made to examine the effect of (+) Gel concentration on injectability of these particle-based formulations (Fig. 8A), which were the injection force, pressure during extrusion, and self-healing behavior. To measure the injection force, an Instron machine was used to extrude particles with 0-3% (+) Gel from a 1 mL needle through a 25G needle at a vertical speed of 0.5 mm/s, corresponding to a volumetric flow rate of 27.54 mL/h (SI Fig. 11). The microgel-based formulations were compared to PBS and a commonly used printing ink, Pluronic (Fig. 8B). The average pressure needed for PBS was 0.18 kPa, and this jumped to ∼4 kPa for 30% Pluronic. The average pressures needed to extrude the particle-based formulations ranged from 0.47 kPa for bare particles to 0.91 kPa for the 1% (+) Gel formulation. The 2% and 3% (+) Gel formulations were within this range at 0.84 and 0.71 kPa respectively. The decrease in force as (+) Gel increases may be due to an increase in repulsion forces as more (+) Gel chains adsorb to the particle surface. No significant differences were found amongst the (+) Gel-containing formulations. The 1% and 2% (+) Gel formulations were significantly different compared to the PBS-based formulation. While the (+) Gel-containing formulations were significantly different compared to the 0% (+) Gel formulation and PBS, these results showed that formulations containing (+) Gel were closer to PBS in terms of injection force compared to 30% Pluronic, illustrating the ease of injecting these (+) Gel-based materials. Significant differences were found between Pluronic and every microgel formulation tested. These results showed that while (+) Gel holds particles together, this type of bonding can be easily broken via a small extrusion force, thus facilitating processes such as injections or 3D printing.

The pressure as a function of total volume extruded (a proxy for time) was measured for each formulation from the injectability tests (Fig. 8C). This data further showed strong similarities in the pressure required to extrude the microgel-based formulations in comparison to PBS. Under the flow conditions used in testing, PBS reached a peak pressure of approximately 0.5 kPa then leveled off after roughly 25 µL had been extruded, maintaining an injection pressure of about 0.18 kPa. Pluronic, in contrast, reached a max pressure of about 4 kPa and leveled off after reaching about 40 µL of ejected volume while maintaining this injection force. The 30% Pluronic, PBS, and 0% (+) Gel formulations showed smooth lines as the ejected volume increased. As (+) Gel is introduced to the microgel-based formulations, there was an increase in pressure fluctuation during ejection. This could be attributed to the presence of aggregate-like clusters, shown in confocal images (Fig. 6A) where pressure would increase with a large cluster, then upon ejection of this cluster, the pressure would decrease.

Rheology was conducted on microgel-based formulations to assess self-healing behavior as a function of (+) Gel concentration over multiple high-low strain cycles (Fig. 8D)^17^. A low oscillatory strain of 1% and high strain of 500% were applied at 1 Hz and were cycled every 60 s for a total of 7 cycles, rapidly inducing transitions that should drive the material to yield from a solid-like state into a liquefied state and back, if the materials are in fact self-healing. All formulations transitioned from solid-like materials to liquid-like behavior as the strain increased, as expected (Fig. 8D). The formulations containing (+) Gel all exhibited self-healing behavior as well as fast recovery as the applied strain was decreased from 500% to 1%. Upon the change in strain, storage moduli values were restored to nearly the original value from the first cycle. This was not observed in the 0% (+) Gel (PBS only) formulation.

Filament formation of each formulation was examined. This was done in air for evaluation, as this can be predictive of the 3D printing process but can also be indicative of use as an injectable hydrogel for non-3D printing applications^17,49–52^. Because the material has poroelastic characteristics, extrusion at high and low flow rates was conducted using a syringe pump to assess any changes in filament formation^44^. The high flow rate was set to 11.7 mL/h, the maximum the pump could achieve with the 1 mL syringe. The low flow rate was set to 1.2 mL/h. The formulations were extruded in 50 µL increments and recorded over time. The maximum filament length prior to filament breakage was measured as a function of (+) Gel over time. The data show that extruded filaments looked nearly identical for all formulations containing (+) Gel, where the filaments were linear, with an aggregate (or bead) of material at the end (Fig. 9A). The 0% (+) Gel formulation, on the other hand, had primarily short filaments, that also had a bead at the end. This was expected, because the (+) Gel tethering of the particles to one another should forces to be sustained along the length of the filament as it begins to extend linearly. In the absence of the electrostatic interparticle interactions, the weight of the bead and filament extruded behind it causes breaks in the filament at shorter lengths (ie. smaller net volumes and mass). The presence of the bead of material at the end of the filaments occurred in all (+) Gel formulations. During extrusion, whose timestamps can be seen in Fig. 9B, the material would kink and roll on itself, followed by ejection of a filament, which did not appear to occur in the PBS-based formulations.

The maximum filament length was measured immediately prior to filament breakage (Fig. 9C). Without (+) Gel, the average maximum filament length observed was approximately 5mm at both high and low flow rates. Upon the addition of (+) Gel, the max filament length increased to roughly 23 mm regardless of (+) Gel concentration. This was also independent of flow rate, as both values for high and low flow rate for each formulation were not statistically significant. To further understand the effect of (+) Gel concentration on ejection of these formulations, all filament lengths were measured and compared (Fig. 9D). The PBS-only formulation had a small distribution of filament lengths. In comparison, the distributions became much larger as (+) Gel was added and did not appear to be dependent on concentration.

Interestingly, during extrusion of the (+) Gel formulations, at both high and low flow rates, the filament would increase in length as expected, but would then retract followed by lengthening then breakage. This can be seen in Fig. 10, where 1% (+) Gel formulations are used as an example. This retraction would sometimes include aggregation of the hanging portion of the filament with the portion emerging from the needle, forming a bead at the filament’s end. This could be attributed to possible relaxation processes within the filament during extrusion. In other words, after the initial bead formed and the filament beg extending, immediately upon leaving the nozzle, extruded material might be in an extended conformation as a function of extrusion rate and nozzle diameter. Electrostatic interactions might then relax this state within the filament, causing a shortening. Continued lengthening afterwards is likely a function of the total weight of the filament dominating opposing forces within the filament that might cause shortening. While this may not impact the material for injectable-based applications, this may impact 3D printed construct shape fidelity.

Because the material compresses during extrusion from a syringe barrel through a needle, confocal images were obtained to assess the change in area fraction and pore size with respect to flow rates (Fig. 11A). As shown in Fig. 11B, area fraction did not change with respect to flow rate or have a trend with (+) Gel concentration. The only statistically significant difference was between the low flow rate and no flow rate groups in the 2% (+) Gel formulation, which showed an increase in area fraction. Pore sizes showed an increase with both high and low flow rates in formulations with (+) Gel. As seen in confocal images (Fig. 11A), the microstructure appeared to change pre- and post-extrusion of (+) Gel-containing formulations, where there appears to be larger pores present. This could be seen in the pore size distributions and average pore size as well (Fig. 11A and 11C), where there was generally an increase in range to include larger pore sizes post-extrusion. This may have implications for cell-biomaterial interactions post-extrusion (in a regenerative setting) or post-printing, as pore size is expected to influence cellular behaviors like migration and proliferation. While there are potential disadvantages if this microarchitectural change affects print fidelity, it is expected that increased porosity might be advantageous in permitting cell behaviors within or on materials, including proliferation and migration.

## Conclusions

Granular formulations of hydrogels are attractive in designing biomaterials that can be delivered by minimally invasive injections in regenerative medicine application or extruded in bioprinting. However, post-deposition stabilization of these materials generally requires that surface-to-surface interactions between microgels within the granular system be designed to prevent further flow or erosion. Here, we have presented an HA-based granular hydrogel that is stabilized through electrostatic interactions with gelatin contained among the particles. This system presents poroelastic, strain stiffening characteristics, which we characterized through rheological analysis and confocal imaging. These properties present options to control porosity within the system. Despite the stabilizing electrostatic interactions, these materials easily flow during extrusion, forming robust filaments as they leave a syringe needle. This work sets the stage for several applications of these materials, such as those in injectable wound healing applications and in 3D printing.

