## Supplemental information for "An injectable granular hydrogel stabilized by electrostatic interactions between hyaluronic acid-based microparticles and soluble gelatin exhibits poroelasticity and strain-stiffening"

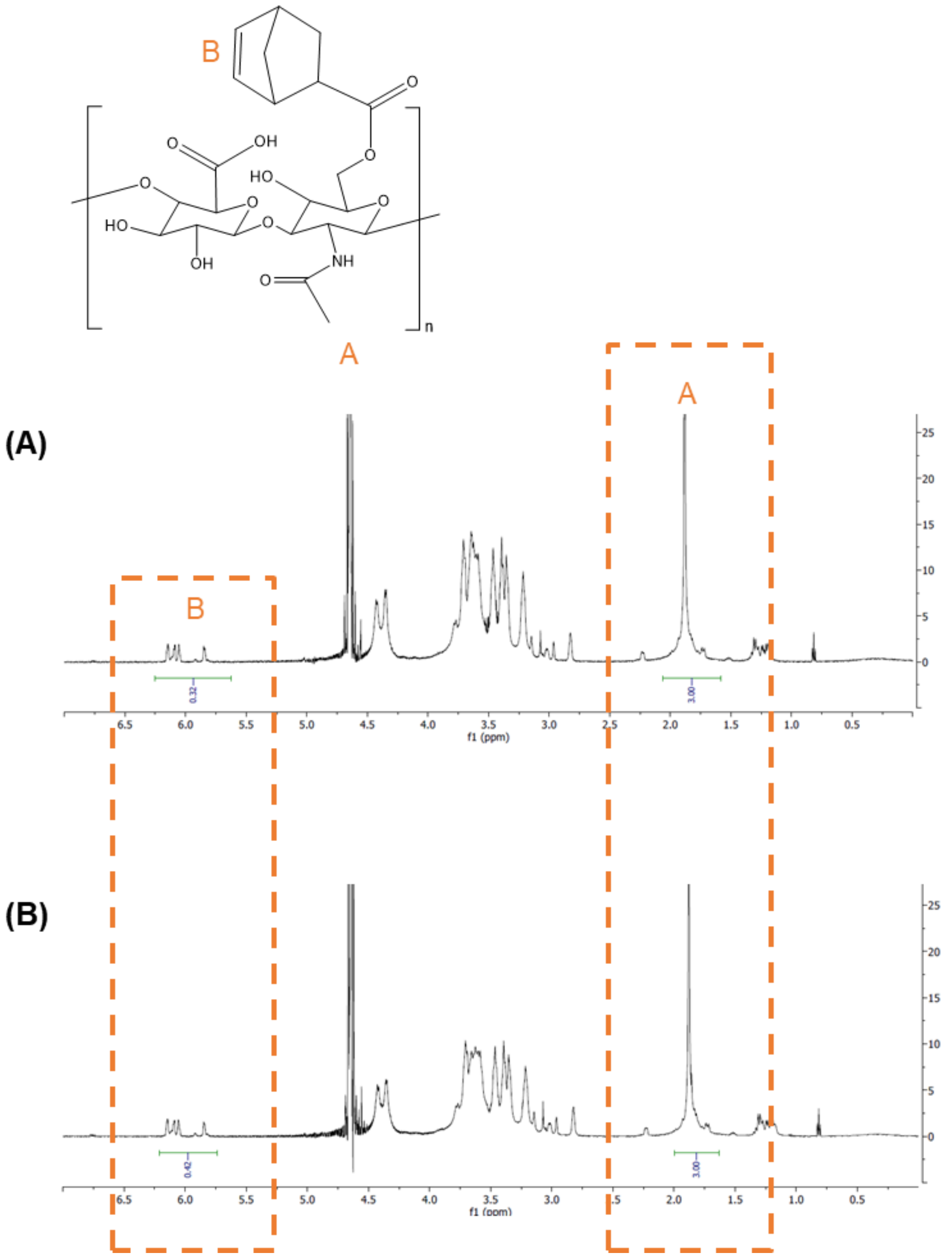


***SI Fig. 1:*** *NMR for two batches of NorHA, one modified to (A) 16% and (B) the other to 21%*

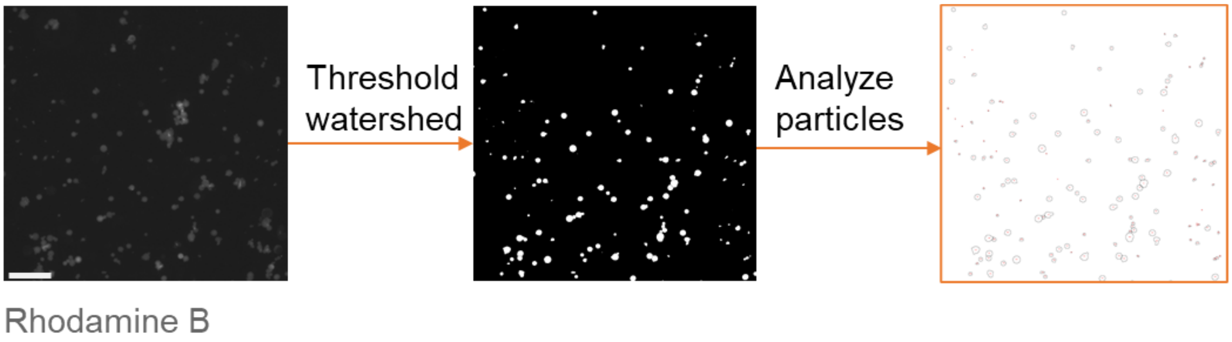


***SI Fig. 2:*** *image processing to obtain NorHA microgel diameters, scalebar=100 µm*

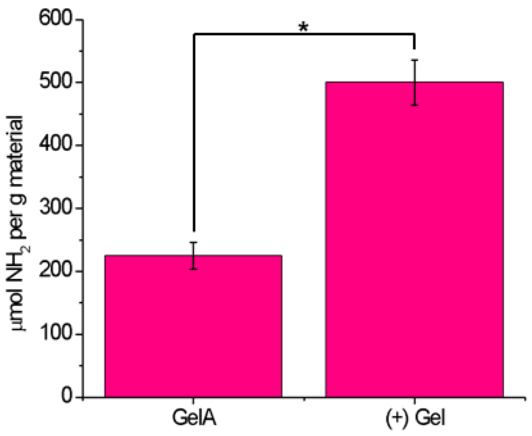


***SI Fig. 3:*** *Fluorescamine on neat gelatin and (+) Gel, * denotes $p<0.05$, error bars are standard deviation, n=3*

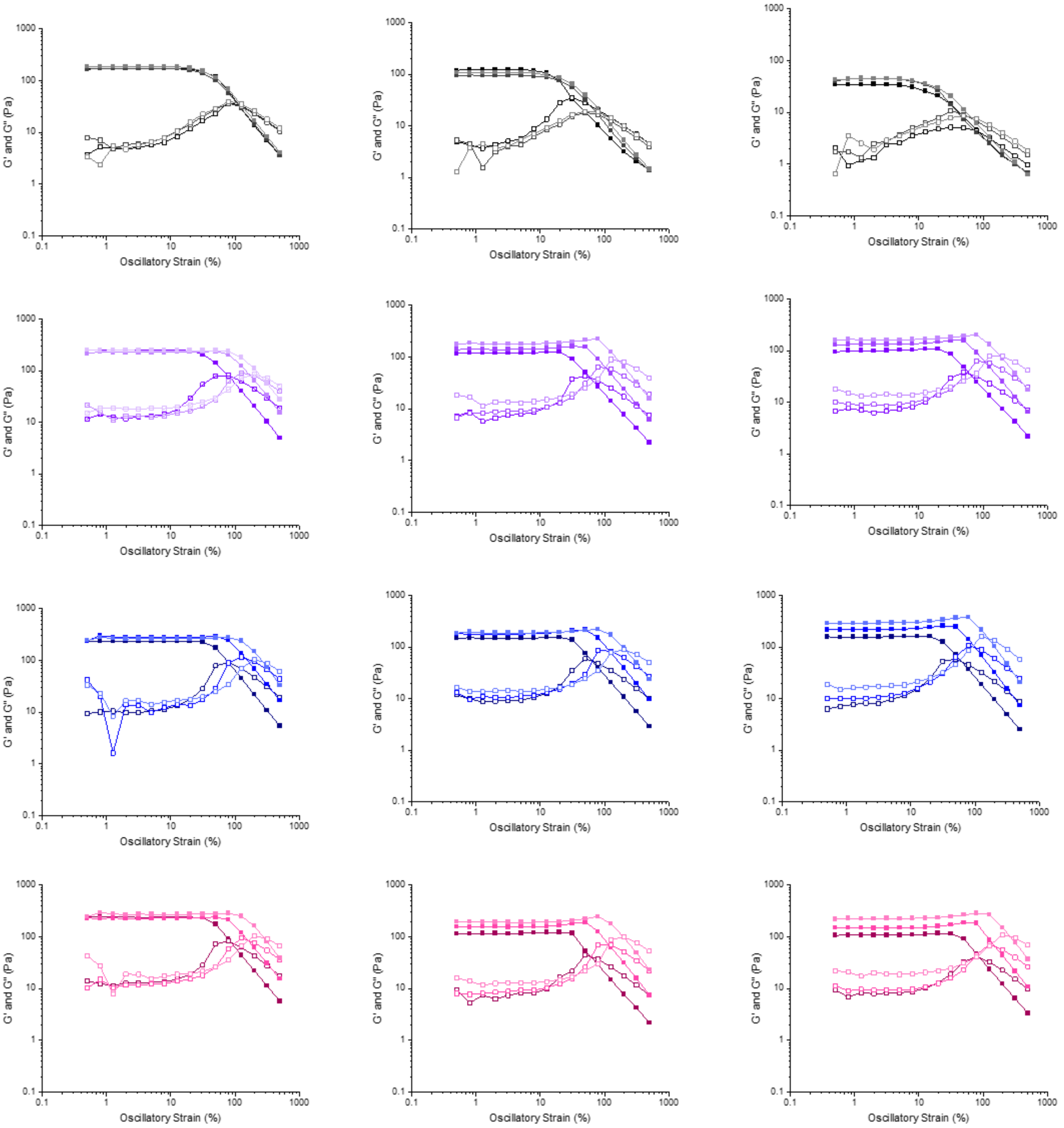


***SI Fig. 4:*** *raw rheological data showing that as compression is increased, particles tethered with (+) Gel show increases in several variables, as analyzed in Fig. 3 of the main text*

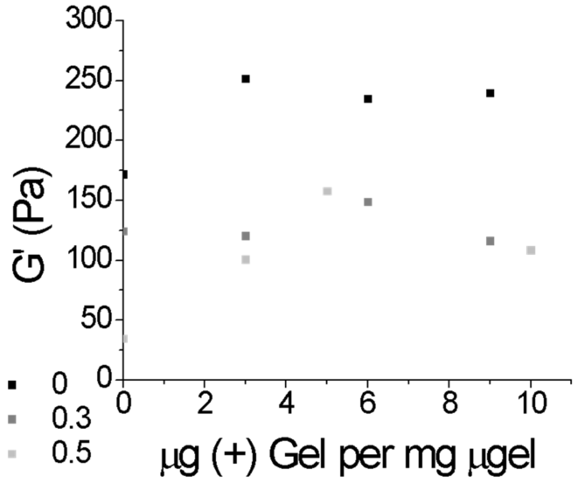


***SI Fig. 5:*** *Dependence of G' on (+) Gel concentration under non-compressive conditions*

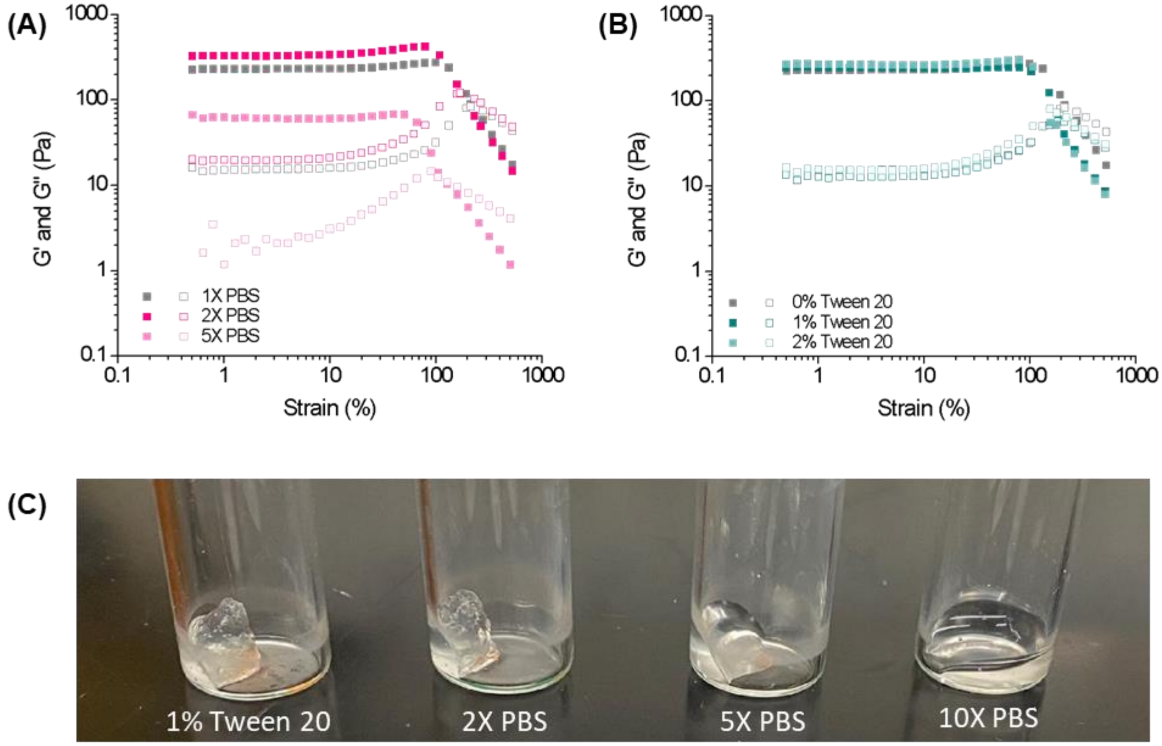


***SI Fig. 6 :*** *Assessment of (A) electrostatic and (B) hydrophobic contributions in the (+) Gel with NorHA microgels, with (C) visual confirmation that the interactions are primarily electrostatic*

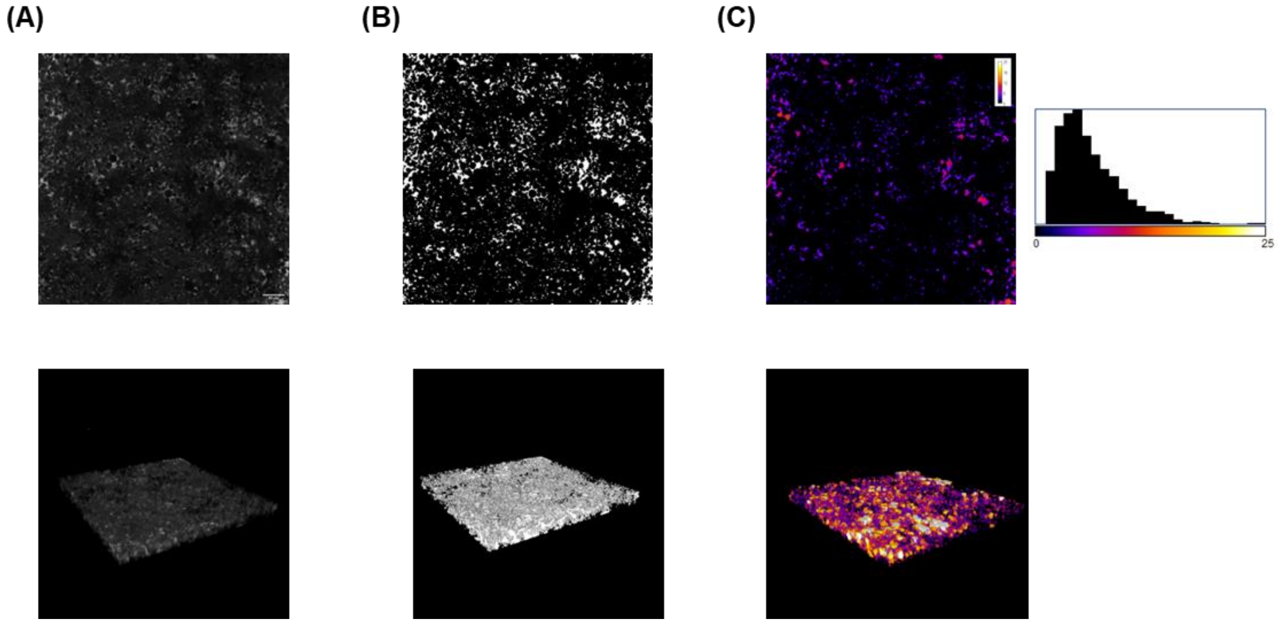


***SI Fig. 7:*** *(A) A confocal image of 2% (+) Gel formulation at a jamming fraction of 0, (B) thresholded image of (A) using Otsu's method, and (C) the trabecular thickness map generated from the thresholded z-stack in BoneJ, from which pore size distributions can be extracted. The second row is a 3D view of the z-stacks in the first row.*

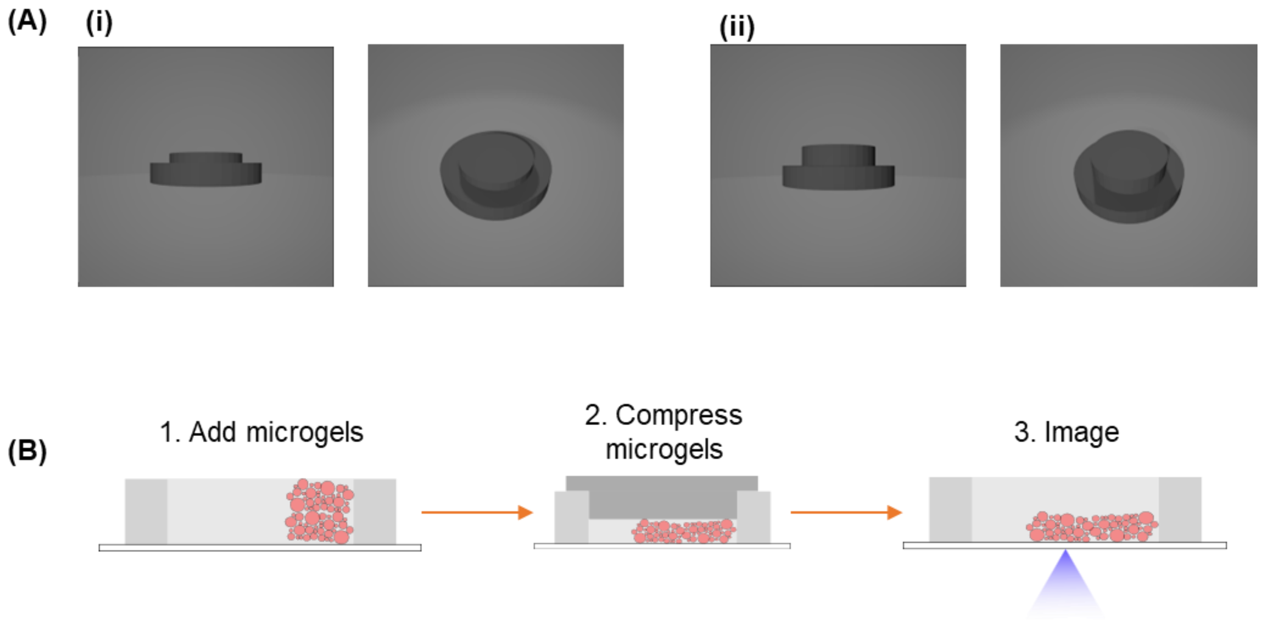


***SI Fig. 8:*** *(A) Compression testing devices for (i) 33% and (ii) 67% compressive strains, and (B) testing setup where material added to PDMS holder, then device used to compress material to selected strain, then imaging done*

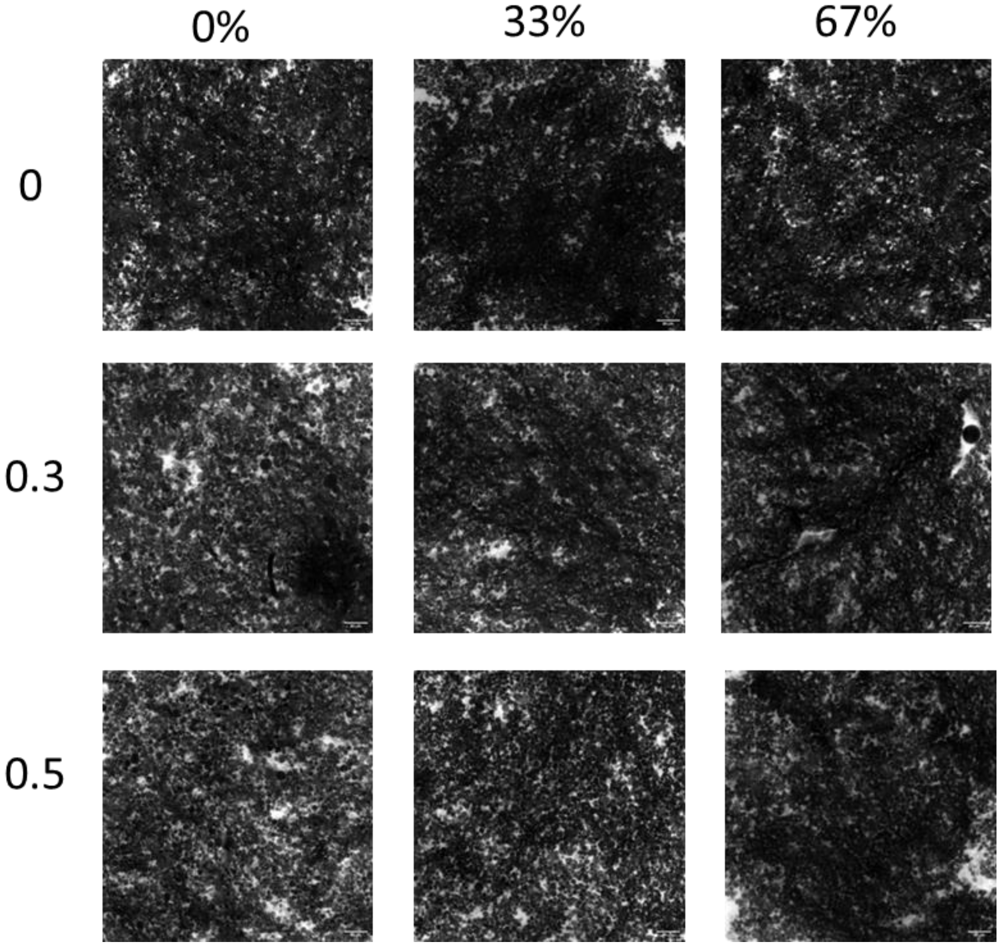


***SI Fig. 9:*** *Confocal images of 2% compression at different compressive strains and jamming fractions*

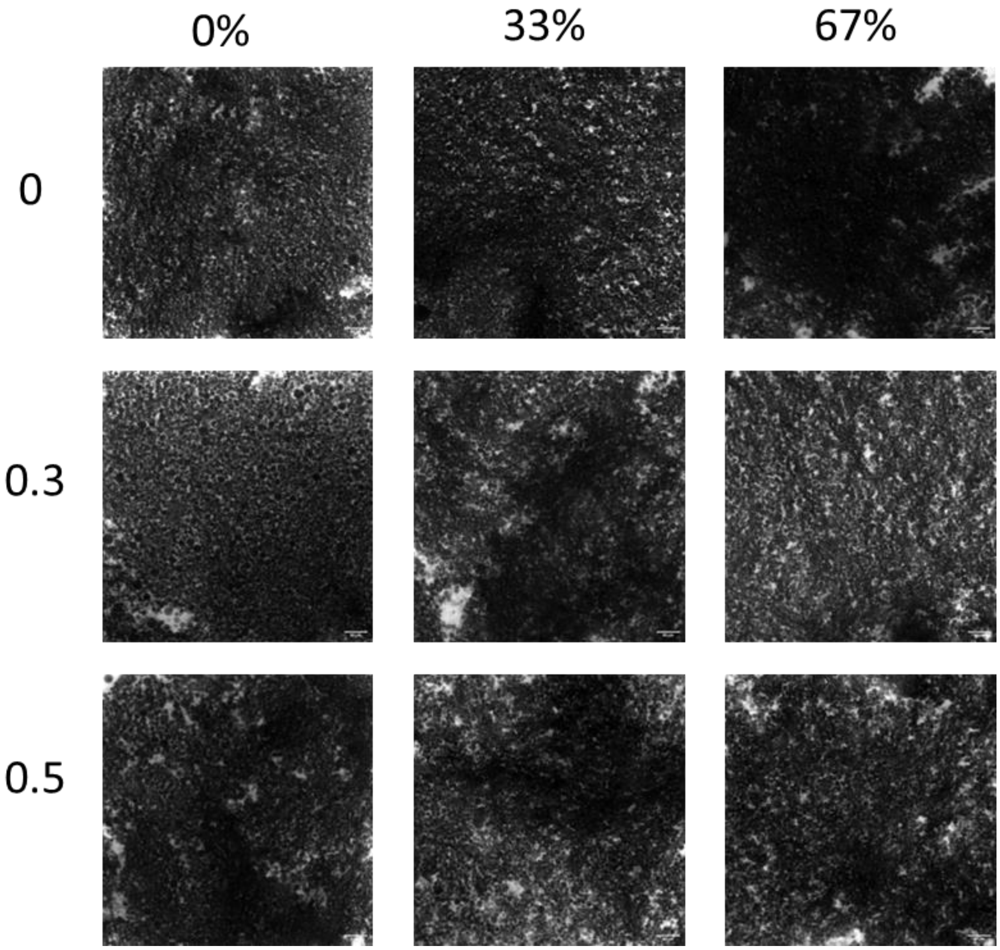


***SI Fig. 10:*** *Confocal 1% (+) gel compression at different compressive strains and jamming fractions*

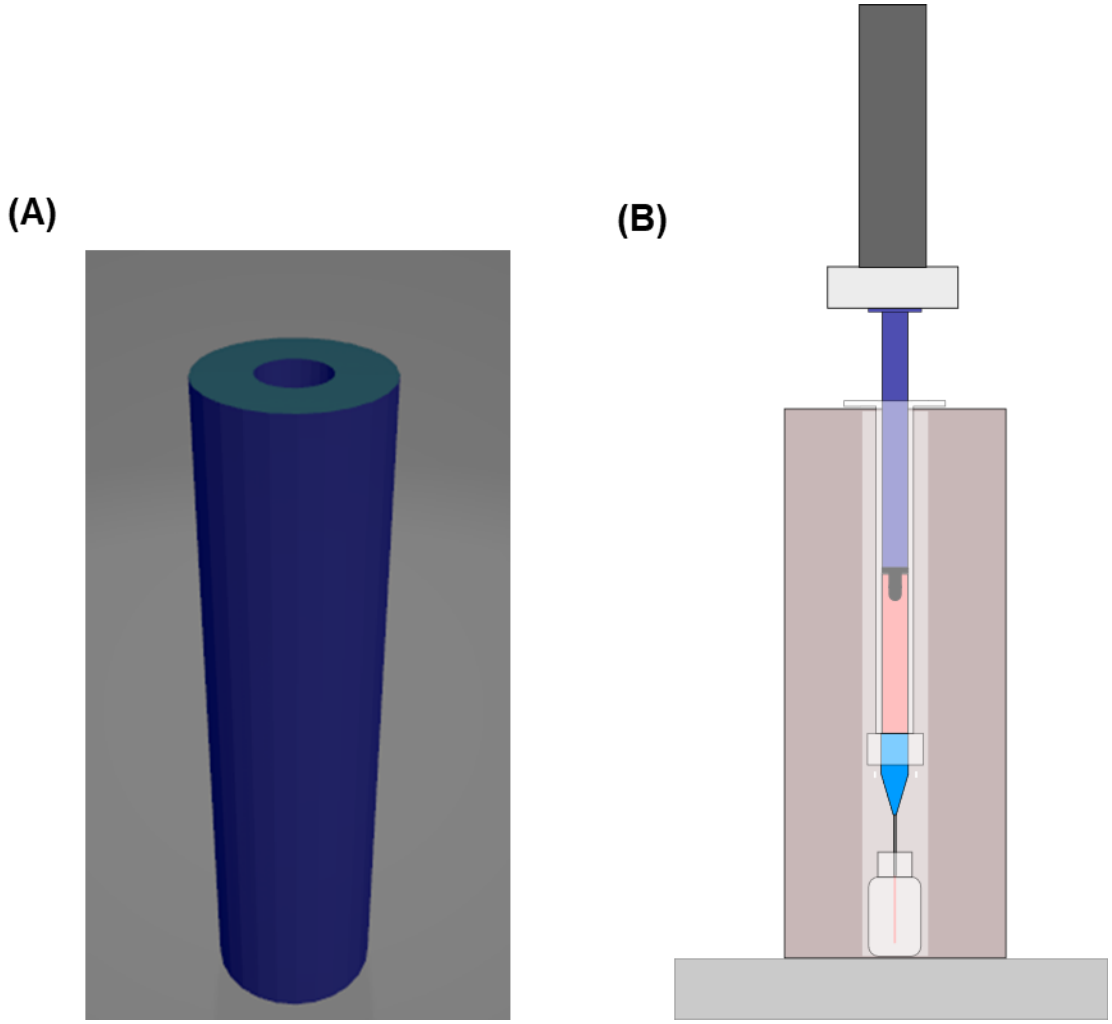


***SI Fig. 11:*** *(A) STL file of syringe holder used on Instron device, and (N) cross-sectional schematic of setup on Instron instrument.*
